# RIC-3 Interacts Directly with the 5-HT_3A_ Receptor to Mediate Trafficking Across Subcellular Compartments

**DOI:** 10.64898/2026.02.15.706004

**Authors:** Nermina Jahovic, Petar N. Grozdanov, Rhea Ramani, Cade Perkins, Clinton C. MacDonald, Joshua Theriot, Quynh Hoa Do, Michaela Jansen

## Abstract

The serotonin type 3A (5-HT_3A_) receptor is a pentameric ligand-gated ion channel (pLGIC) in central and peripheral neurons that conducts sodium and potassium ions upon serotonin binding. 5-HT3 receptors modulate neurotransmission and synaptic plasticity, influencing mood, sleep, appetite, and addiction. Disruptions in serotonin signaling are linked to central nervous system disorders, including schizophrenia, anxiety and depression. Clinically, these receptors are targeted by antagonists to treat chemotherapy-induced nausea and vomiting. The functional surface expression of these channels is regulated by the chaperone protein Resistant to Inhibitors of Cholinesterase 3 (RIC-3) that promotes plasma-membrane expression, maturation, and trafficking of 5-HT_3A_ and nicotinic acetylcholine receptors. Our previous work identified a duplicated RIC-3 binding motif within the 5-HT_3A_ intracellular domain (ICD). However, it was unclear whether this interaction reflected native conditions. Here, we used a recombinant 5-HT_3A_ ICD peptide in peptide-resin pull-down assays to investigate RIC-3 Interactions in plasma membrane (PM) fractions from *Xenopus* oocytes, endoplasmic reticulum (ER) fractions from SH-SY5Y cells, and mouse brain tissue. Across all tested systems, the 5-HT_3A_ ICD peptide specifically bound RIC-3. Furthermore, RIC-3 knockdown (RIC-3 KD) SH-SY5Y cells showed a marked reduction in peptide binding and decreased surface levels of nAChRα7 and 5-HT_3A_ receptors. These results demonstrate RIC-3-5-HT_3A_ ICD interaction in native cellular contexts and support a role for RIC-3 in regulating receptor surface expression and neuronal signaling.

## 1. Introduction

Serotonin 5-HT_3_ receptors mediate rapid excitatory signaling in the central and peripheral nervous systems. In the gastrointestinal system, which produces ∼95% of the body’s serotonin, these receptors regulate gut motility, sensory signaling, and nausea, making them essential for normal digestive physiology and important therapeutic targets for gastrointestinal disorders.

5-HT_3A_ receptors are members of the pentameric ligand-gated ion channel (pLGIC) or Cys-loop receptor family, characterized by a conserved 13-amino acid disulfide-linked loop in the N-terminal extracellular domain. This family also includes nicotinic acetylcholine receptors (nAChRs), gamma-aminobutyric acid receptors (GABA-ARs), and glycine receptors (1). Dysregulation across this receptor family, including 5HT_3A_ receptors, has been linked to major psychiatric conditions such as anxiety, depression, schizophrenia, addiction and autism-spectrum traits (2) (3) (4).

The 5-HT_3A_ receptor is integrated into the cell membrane as a homomeric assembly of five subunits that encircle a central pore (**Fig. 1)**. Each subunit contains an extracellular N-terminal domain (ECD) with agonist-binding sites, a transmembrane domain (TMD) with four α−helical segments (M1–M4), and an intracellular domain (ICD) critical for channel conductance and regulation (5) (6). Disulfide-crosslinking studies of the ICD have shown that changes to the relative distance/orientation of the upper but not the lower part of the membrane associated (MA) helix are required for ion conduction (7). While ECD and TMD have been well characterized by X-ray crystallography and cryo-electron microscopy (8) (9) (10) (11) (12) the ICD has been less studied. Recent NMR and ESR studies revealed a highly flexible α7 nAChR ICD (13) in addition to the MA-helices previously resolved for 5-HT_3A_ receptors (10) (8). The ICD of pLGICs serves as a target for post-translational modifications including palmitoylation, phosphorylation, and ubiquitination that profoundly impact receptor function (14) (15) (16) (17) (3) (18) (19).

**Fig. 1.**
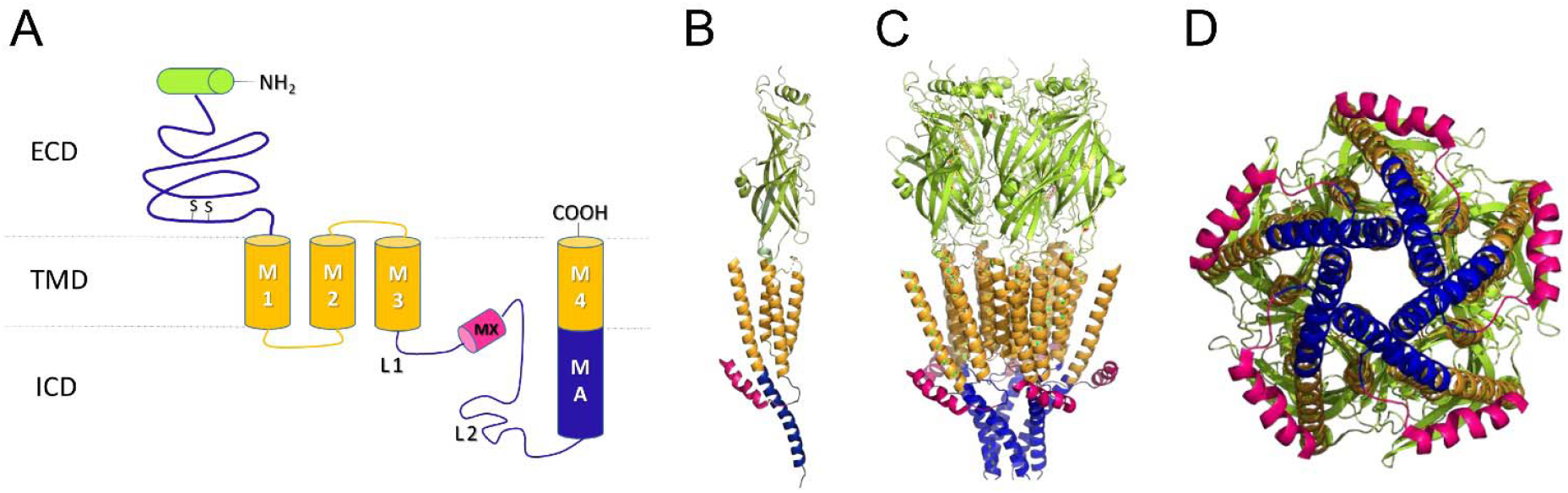
Structure of the 5-HT_3A_ receptor. **A** Scheme of a single subunit demonstrating the arrangement of the three domains, extracellular (ECD), transmembrane (TMD) with four α-helical segments (M1 through M4; yellow), and intracellular (ICD blue and magenta) domain with a 17 amino acid long MX-helix and a MA helix of 31 amino acids continuous with the subsequent M4 transmembrane helix. **B** and **C** Ribbon representation of single subunit and pentameric receptor, respectively. The views are parallel to the membrane plane and colors are as in panel A. **D** Pentameric receptor as viewed from the intracellular side. (PDB ID 4PIR; Hassaine et al., 2014).

Resistant to Inhibitors of Cholinesterase 3 (RIC-3) is an endoplasmic reticulum (ER) membrane protein with chaperone activity, facilitating assembly of nAChRs (particularly the α7 subtype) and 5-HT_3_ receptors (20). Initially identified in *Caenorhabditis elegans*, RIC-3 enhances nAChR surface expression (15) (21). Moderate increases in RIC-3 expression facilitate receptor assembly, whereas excessive RIC-3 expression caused ER retention of nAChRs, potentially due to the RIC-3 oligomerization through its cytosolic coiled-coil domain (22).

Previous work demonstrated that replacement of the large intracellular M3M4 loop of 5-HT_3A_ and GABA-A-ρ receptors with the prokaryotic heptapeptide (SQPARAA) linker, generated functional receptors (6) but abolished RIC-3-dependent enhancement of 5-HT_3A_ surface expression in *Xenopus laevis* oocytes. Conversely, insertion of the nAChR-α7 or 5-HT_3A_ ICD into the prokaryotic *Gloeobacter violaceus* (GLIC) channel conferred RIC-3 sensitivity (23) (24), emphasizing the ICD’s requirement for RIC-3 mediated regulation. RIC-3 interacts with 5-HT_3A_ receptors during biogenesis, modulating their maturation and functional responsiveness to serotonin (25) (26) (27). Sequential 5-HT_3A_ ICD deletions identified a 24-amino-acid L1-MX-segment within the 115-residue 5-HT_3A_-ICD as sufficient for RIC-3 binding (28). This interaction was further validated using alanine-scanning, pull-down assays, and two-electrode voltage-clamp (TEVC) recordings (25). Further truncation of this segment to create the MX-peptide was sufficient for interaction with RIC-3. Our studies revealed a duplicated DWLR…VLDR motif, in the MX-helix and MA-M4 junction, that is essential for RIC-3 recognition.

In the present study, we used the L1-MX peptide covalently coupled to resin to probe its interaction with RIC-3 from native sources, specifically mammalian ER fractions prepared from SH-SY5Y neuroblastoma cells and mouse brain tissue and compared these results with *Xenopus* oocytes heterologously expressing RIC-3 as a positive control. Our hypothesis was that L1-MX will pull down RIC-3 from native ER fractions similarly to heterologously expressed RIC-3, indicating a bona fide interaction in vivo.

## 2. Materials and methods

### 2.1 Injection and expression of RIC-3 protein in *Xenopus laevis* oocytes

*Xenopus laevis* oocytes were purchased from a licensed commercial supplier, Ecocyte Bioscience US LLC (Austin, TX). Oocyte preparation procedures and RIC-3 cRNA injection and expression were performed as previously described (25).

### 2.2 Mouse brain cell fractionation

All animal treatments and tissues obtained in the study were performed according to protocols approved by the Institutional Animal Care and Use Committee at the Texas Tech University Health Sciences Center in accordance with the National Institutes of Health animal welfare guidelines. TTUHSC’s vivarium is AAALAC-certified and has a 12/12-hour light/dark cycle with temperature and relative humidity of 20–22 °C and 30–50%, respectively.

Mouse brain lysis was performed using a protocol adapted from procedures established for rat liver (29) (**Fig. 2A**). Only essential modifications are described here. Brains from 2–5 C57BL/6NCrl (Charles River) mice (∼4 months old, male/female siblings; average brain weight ∼0.5g each) were extracted, homogenized, and stored at −80 °C until use.

**Fig. 2.**
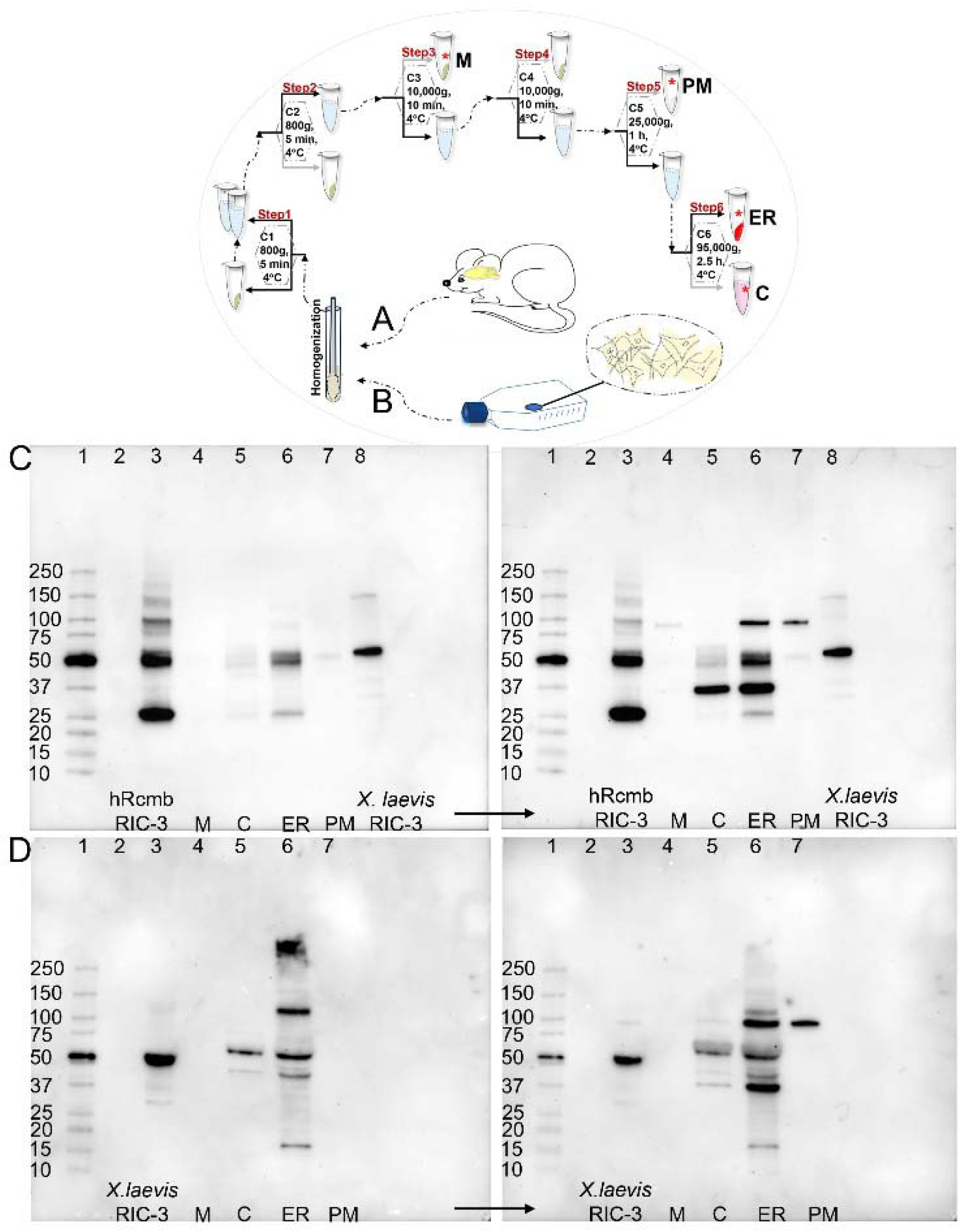
Subcellular fractionation of mouse brain tissue or SH-SY5Y cells. Illustration of the fractionation procedure for isolating crude plasma membrane (PM) and endoplasmic reticulum (ER) fractions from **A** mouse brain or **B** SH-SY5Y neuroblastoma cells. Additionally, cytosolic (C), and crude mitochondrial (M) fractions are obtained. Red star denotes samples used for subsequent analyses. C1-C6 depicts differential centrifugations. **C** Subcellular localization of RIC-3 in mouse brain and SH-SY5Y cells; validation of fraction purity. RIC-3 protein was probed in cytosol (C), endoplasmic reticulum (ER), mitochondria (M), and crude plasma membrane (PM) fractions using a mouse anti-RIC-3 primary antibody (Abnova, cat# H00079608-B01P) and an HRP-conjugated goat-raised anti-mouse secondary antibody (Thermo Fisher Scientific, Cat. #31430). Analyses were performed using fractionated mouse brain tissue **C** and SH-SY5Y cells **D**. Human recombinant RIC-3 (hRIC-3) and RIC-3 (both ∼50 kDa) expressed in *Xenopus laevis* oocytes were included as positive controls. In both systems, the strongest RIC-3 (∼50 kDa) signal was observed in the ER fraction. In mouse ER fractions (panel C), RIC-3 appears as a doublet at ∼50 kDa (lane 6), consistent with post-translational modification. Fraction purity was validated using calnexin (ER; Invitrogen cat# PA5-34754; ∼100 kDa), Na^+^/K^+^-ATPase (PM; Cell Signaling Technologies cat# 3010S; ∼100 kDa) and GADPH (C; Cell Signaling Technologies, 5174; ∼35 kDa). HRP-conjugated goat anti-rabbit secondary antibody (Thermo Fisher Scientific, cat# 31460) was used for rabbit-raised primaries. Blots shown in panels **C** and **D** (to the right of the arrow) were performed sequentially on separate days. Blots are semiquantitative, and ER signal bleed-through in non-ER fractions was corrected using a Calnexin-based linear contamination model. Additional RIC-3 reactive bands in mouse brain/SH-SY5Y cells correspond in size to mouse and human RIC-3 isoforms in the Uniprot database.

### 2.3 SH-SY5Y cell line fractionation

SH-SY5Y cells (Sigma Aldrich, cat. #94030304) were cultured in DMEM/F12 Complete medium supplemented with 2 mM L-glutamine and 10% FBS (Sigma Aldrich cat. #SLM-243-B), and 1% Penicillin-Streptomycin (5,000 U/mL; Life Technologies). Cells were maintained at 37 °C in 5% CO_2_ and 95% air. Passages 3–7 were selected for the fractionation process. When the cells reached approximately 85– 90% confluence, they were collected using a scraper (Falcon cat. #353085). SH-SY5Y cells were routinely passaged with trypsin; however, for experimental harvesting we used cell scraper instead, since Western blot analysis showed that exposure to trypsin during harvesting led to RIC-3 degradation producing smeared and degraded bands.

We fractionated SH-SY5Y cell lysates (**Fig. 2B**) as for mouse brain cells but minimized the solubilization amount to the smallest feasible quantity (200–500 μl per fraction) as determined through visual assessment of the pellet yield to be dissolved and validated by Western blot analysis of samples from both mouse and SH-SY5Y cells. The sample loading amount (approximately 10 μg of total protein) for each Western blot was determined by measuring the sample concentrations using the BCA kit (Micro BCA Protein Assay Kit, Thermo Fisher Scientific, Cat. No. 23235).

### 2.4 RIC-3 probing and fraction verification through SDS-PAGE/immunoblotting

Each flow-through sample was analyzed by SDS-PAGE and Western blotting using the Bio-Rad Trans-Blot Turbo system, following the manufacturer’s instructions.

Following transfer, membranes were blocked for 1 hour at RT in 5% nonfat milk prepared in TTBS (0.1% Tween-20, 100 mM Tris, 0.9% NaCl, pH 7.5) and incubated for 2 hours at 4 °C under gentle agitation. Blots were probed overnight at 4 °C with mouse anti–RIC-3 antibody (Abnova, #H00079608-B01P, 1:2,000). After three 5-minute washes with TTBS, membranes were incubated for 2 hours with HRP-conjugated goat anti-mouse secondary antibody (Thermo Fisher, #31430, 1:5,000), washed again three times, and developed with SuperSignal West Femto substrate (Thermo Fisher). Protein bands were visualized using a ChemiDoc MP Imaging System (Bio-Rad).

To assess subcellular fraction purity, membranes were probed with antibodies against housekeeping proteins: calnexin (ER marker, Invitrogen, #PA5-34754), Na^+^/K^+^-ATPase (plasma membrane marker, Cell Signaling Technologies, #3010S), and GAPDH (cytosolic marker, Cell Signaling Technologies, #5174).

Western blot bands were quantified using Bio-Rad ImageLab software, and all intensities were normalized to total protein so that differences reflected true variations in protein expression across samples. Calnexin, an ER-resident chaperone, was used as both an internal control (**Fig. 3B**) and a marker of ER contamination (**Fig. 2C-D**). For each non-ER fraction X (mitochondria, cytosol, PM), the ER contamination factor was calculated as:

**Fig. 3.**
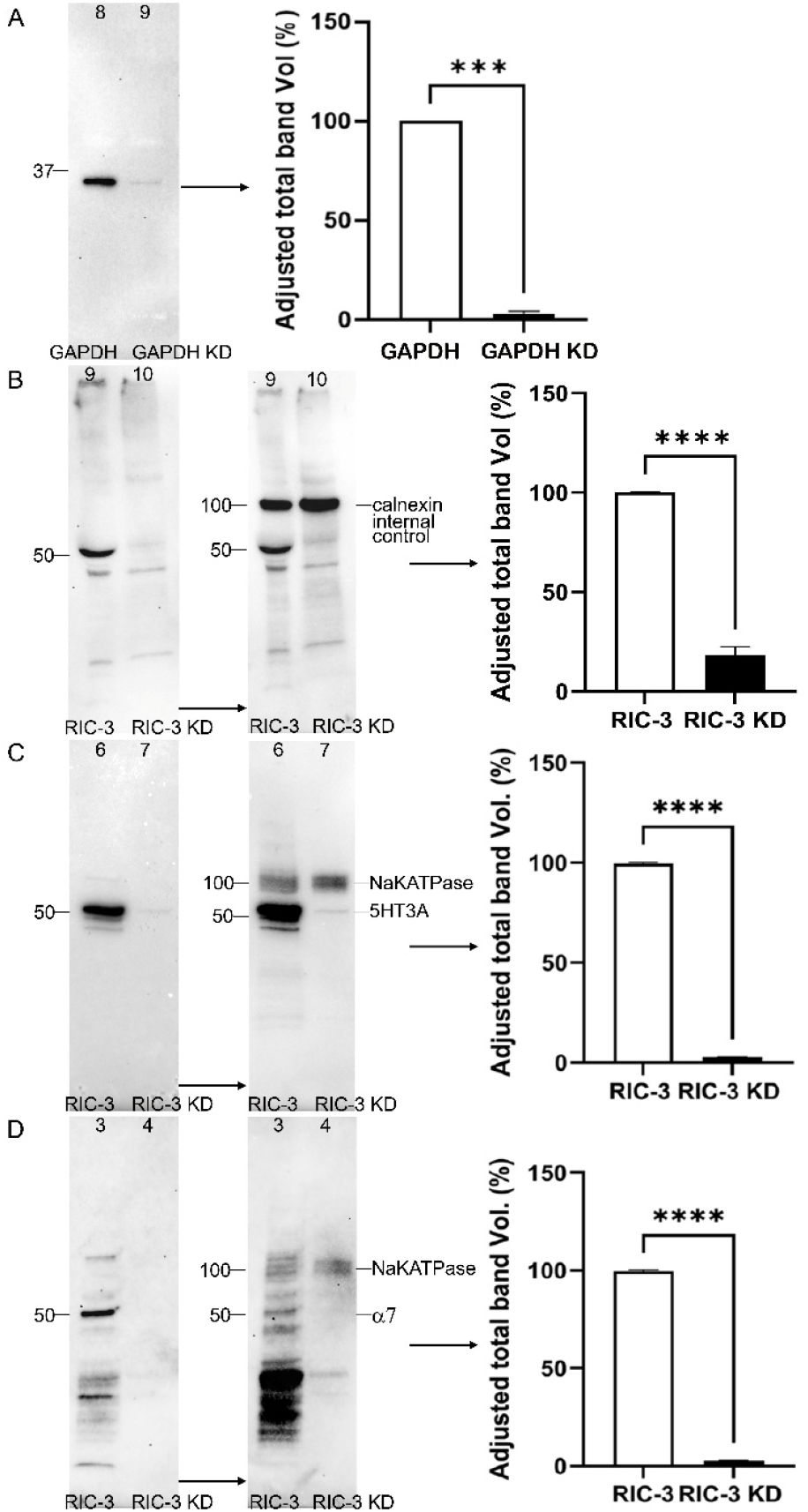
RIC-3 knockdown in SH-SY5Y cells. **A** For control experiments, SH-SY5Y cells treated with 5 nM Silencer GAPDH siRNA (ThermoFisher Scientific, Cat# 4390849), achieved approximately 95% GAPDH knockdown (KD). The cells were incubated for 72 hours at a confluence level of 50%. **B** RIC-3 protein knockdown was performed by incubating SH-SY5Y cells at 50% confluence with 5 nM RIC-3 siRNA for 24 hours. The first Western blot (left) shows the control (RIC-3; 50 kDa, lane 9) and siRNA-treated samples for RIC-3 (RIC-3 KD; lane 10). To validate the knockdown, we used calnexin (100 kDa; lane 9-10) as an internal loading control by probing the blot shown on the left with calnexin antibody to yield the blot on the right. **C-D** RIC-3 knockdown reduces plasma membrane surface expression of nACh α7 and 5HT3A receptors in SH-SY5Y cells. Western blot analysis was performed on plasma membrane (PM) fractions isolated from SH-SY5Y cells either transfected with RIC-3 siRNA (RIC-3 KD) or untransfected (RIC-3). Blots were probed with antibodies against **C** 5-HT3A (50 kDa) and **D** nAChRα7 (50kDa) subunits. Both showed approximately 98% reduction in the PM fraction of RIC-3 knockdown cells compared to control. Na^+^/K^+^-ATPase (100 kDa) was used as a reference protein for sample loading normalization. All blots were quantified and normalized using Bio-Rad ImageLab software. Blots in Fig. 3 were cropped for clarity and presentation purposes. Molecular-weight marker positions (in kDa) are indicated on the sides of each blot.

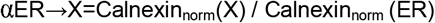

where αER→XαER→X represents the fraction of ER signal present in fraction X. To generate **Fig. 2C-D**, the Calnexin signal in each non-ER fraction was then corrected by subtracting this ER contribution:

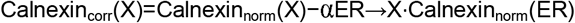

If subtraction resulted in a negative value, it was interpreted as complete ER bleed-through, and the corrected intensity was set to zero. Calnexin bleed-through correction was required only for the mitochondria and PM fractions (**Fig. 2C-D**). A similar contamination-correction approach has been used to quantify subcellular compartment-specific signals by adjusting for fraction bleed-through using organelle markers and linear deconvolution (30).

### 2.5. RIC-3 knockdown in SH-SY5Y cells

RIC-3 knockdown was achieved through transient transfection using Lipofectamine RNAiMAX (Thermo Fisher, Cat. #13778150). SH-SY5Y cells were transfected at 50% confluence with 5□nM siRNA targeting RIC-3 (Thermo Fisher). GAPDH siRNA (Silencer Select, Cat. #4390849, Thermo Fisher) was used as positive control. Each siRNA was diluted in OptiMEM (Thermo Fisher, Cat. #11058021) at a 1:100 ratio and mixed with Lipofectamine at a 1:1 ratio before being added to cells.

### 2.6 Pull down assay

#### 5-HT_3_ receptor ICD peptide (L1-MX)

The preparation of the 5-HT_3A_ ICD L1-MX-peptide, its covalent coupling to iodoacetyl resin, and the subsequent pull-down assays with RIC-3 were performed as previously described (25) (**Fig. 4A**). Only experimental conditions specific to this study are noted.

**Fig. 4.**
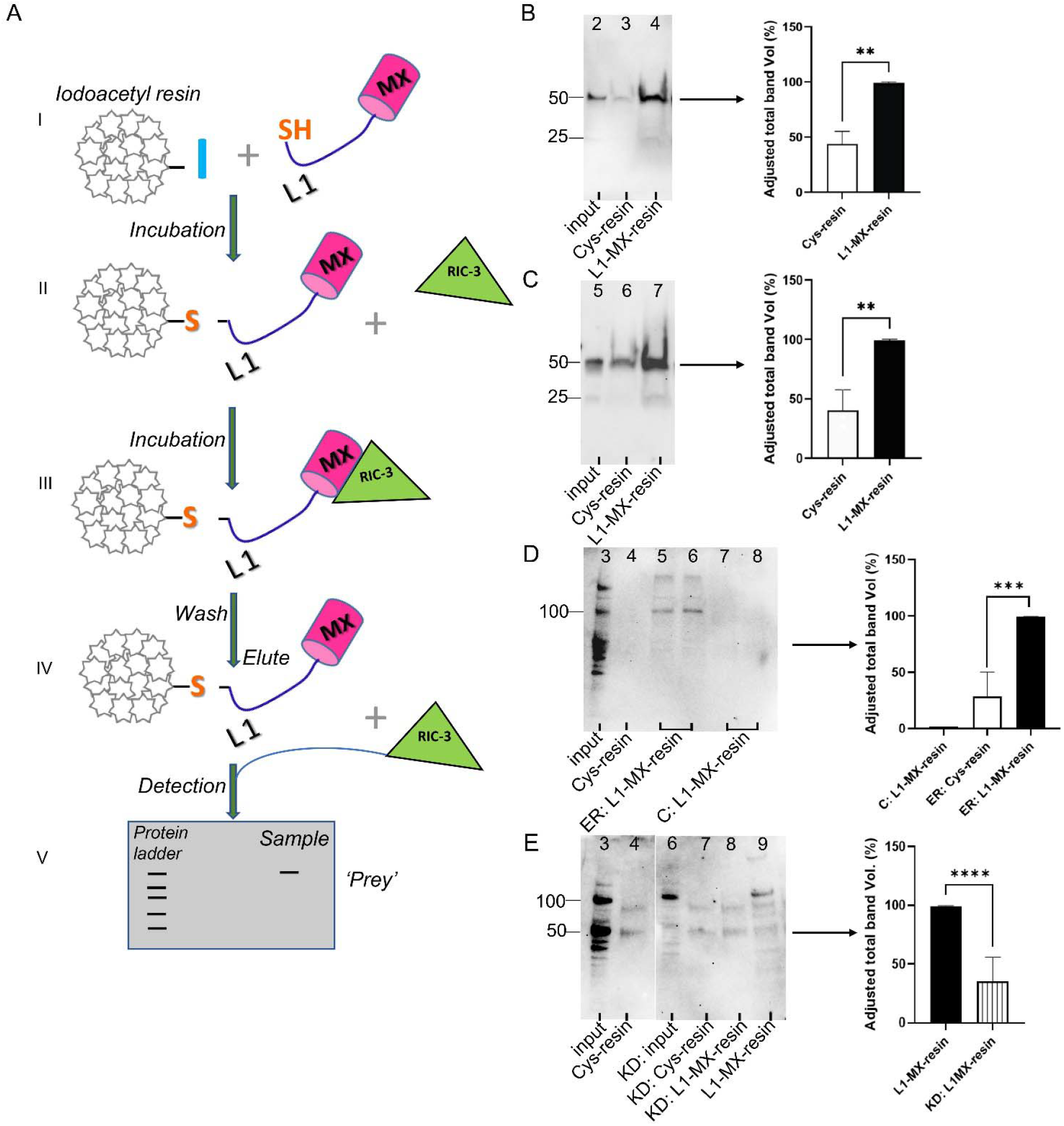
Identification of RIC-3 following affinity enrichment using immobilized L1-MX peptide. **A** Pull-down assay procedure. **I** L1-MX peptide containing an N- or C-terminal Cysteine (‘bait’) and the iodoacetyl resin react to form a covalent bond yielding L1-MX peptide alkylated with resin as shown in **II** during the coupling process. **III** Tissue or cell extract containing RIC-3 protein are incubated with the modified resin and RIC-3 will bind to the linked L1-MX peptide. **IV** Elution with SDS denatures this complex to release bound proteins/RIC-3 that had interacted with L1-MX peptide. **V** Eluates are separated by denaturing SDS-PAGE and RIC-3 detected using immunoblotting. **B-E** Pull-down results for **B** *Xenopus laevis* oocytes PM, **C** mouse brain ER, **D** SHSY5Y ER and cytosol (C) fractions and **E** ER fractions from transfected SH-SY5Y cells treated with 5□nM RIC-3 siRNA. Pull-downs from oocytes **B** and mouse brain **C** confirmed RIC-3 binding to L1MX (50 kDa; lane 4 and 7, respectively). In SHSY5Y cells, cytosol lysates **D** (lane 7-8) did not recover RIC-3, indicating that enrichment was specific to the ER fraction. ER-derived RIC-3 (100 kDa; lane 5-6) bound ∼3-fold more strongly to L1-MX than to the Cys-resin control **D.** In panel **E**, RIC-3 interacts with L1-MX in non-transfected cells (100 kDa band, lane 9), but not in RIC-3 knockdown samples (lane 8). Faint additional bands at ∼50 and ∼75 kDa (lanes 4, 6-9) represent nonspecific signals. Blots in panels B-E were cropped for clarity and presentation purposes. Molecular-weight marker positions (kDa) are indicated on the sides of each blot.

#### L1-MX-iodoacetyl resin coupling

The peptide was coupled to the resin to achieve a resin-bound peptide/resin ratio of 5.19 to 9.13 mg/ml which was determined and optimized for each coupling reaction. The coupling reaction was carried out overnight at 4 °C with gentle rotation. Free reactive sites on the resin were blocked with L-cysteine (Cys-capped resin) by incubating the resin overnight at 4 °C with gentle shaking (25).

#### RIC-3 pull-down

Pull-down assays using mouse brain- and oocyte-derived samples with *L1-MX-* resin to isolate RIC-3 were performed as previously described (25) For SH-SY5Y samples, binding reactions were carried out overnight at 4 °C with gentle rotation. Proteins were eluted using a gentle buffer containing 50□mM Tris-HCl (pH□7.4), 150□mM NaCl, 10% glycerol, 0.01% digitonin, 0.3□mM TCEP, and Laemmli buffer. Samples were incubated at 50 °C for 5 minutes, then centrifuged at 1,000 ×*g* for 2 minutes at RT.

### 2.7. Deglycosylation of native glycoproteins

To assess N-linked glycosylation under native conditions, glycoprotein samples were treated with PNGase F (NEB #P0708S) or Endo H (NEB #P0702S) following the manufacturer’s non-denaturing protocols.

### 2.8. α7 and 5-HT_3A_ surface expression following RIC-3 knockdown in SH-SY5Y cells

Western blot analysis was performed on PM fractions isolated from SH-SY5Y cells transfected with RIC-3 siRNA (RIC-3 KD) or untransfected (RIC-3). Membranes were probed with antibodies against the α7 nAChR subunit (Santa Cruz Biotechnology, Cat. No. sc-58607) and the 5-HT_3A_ receptor (GeneTex, Cat. No. GTX13897). Na^+^/K^+^-ATPase was used as a reference protein for sample loading normalization.

## 3. Results

In our previous work, we identified a RIC-3 binding motif within the MX and MAM4 segments of the 5-HT_3A_ ICD, using recombinant RIC-3 from *E. coli* or *Xenopus laevis* oocytes (26) (27) (23) (6). Given the role of 5-HT_3A_ receptors in neurological and psychiatric disorders, we asked whether the 5-HT_3A_ L1-MX peptide also binds RIC-3 natively expressed in human cells and mouse brain. To address this, we isolated RIC-3 containing fractions from mouse brain and SH-SY5Y cells and assessed their binding to immobilized 5HT_3A_ L1-MX peptide (**Fig. 2** and **4**).

### 3.1 Fraction purity validation and subcellular localization of RIC-3

Fresh mouse brain tissue was fractionated to isolate crude mitochondria, cytosol, ER, and crude PM (29) (31) (25). The same protocol was applied to SH-SY5Y neuroblastoma cells (**Fig. 2A-B**), enabling collection of all four fractions from a single preparation, an advantage over single-fraction kits (32). Calnexin, a chaperone predominantly localized to the ER, validated ER fractions, while Na^+^/K^+^-ATPase marked the PM. Western blot analysis confirmed enrichment in both mouse brain and SH-SY5Y fractions, with strong bands observed at ∼100 kDa for calnexin and Na^+^/K^+^-ATPase, (**Fig. 2C-D**). The cytosolic fraction was validated with GAPDH (∼35 kDa) (Cell Signaling Technologies, cat. no. 5174). PM fractions were prepared, but due to low protein yield they were excluded from the pull-down assays.

RIC-3 (∼50 kDa) was readily detected in mouse brain tissue/SH-SY5Y cell line ER fractions (**Fig. 2C-D**). To account for ER carryover into non-ER fractions, we calculated a calnexin-derived contamination factor (αER→XαER→X) and subtracted from the apparent signal. This correction eliminated ER bleed-through, allowing a more accurate representation of protein distribution across compartments. The correction was applied only to calnexin, while experimental quantification of RIC-3 relied on standard normalization to the appropriate compartment-specific markers (**Fig. 2C-D**).

### 3.2 RIC-3 protein in *X. laevis*, mouse brain and human SH-SY5Y cell line

For comparison, human recombinant RIC-3 (Novus Biologicals, cat# NBP238058PEP) and RIC-3 overexpressed in *X. laevis* oocytes were included. As reported previously (25), a distinct ∼50□kDa band was observed in RIC-3-injected oocytes, consistent with the current study (**Fig. 2C-D**).

Using anti-RIC-3 antibody, endogenous RIC-3 was detected in native ER fractions. Mouse ER fractions showed ∼50 kDa and ∼25 kDa bands (**Fig. 2C**, lane 6; blot to the left), while SH-SY5Y ER fractions displayed ∼100, 50, 37, and 15 kDa bands (**Fig. 2D**, lane 6; blot to the left), corresponding to known human (Q7Z5B4) and mouse (Q8BPM6) isoforms. Given that the predicted molecular weight (MW) of RIC-3 is ∼44 kDa, the slower-migrating ∼50□kDa band likely reflects N-linked glycosylation, consistent with prior reports of glycosylated RIC-3 in mammalian systems (33).

### 3.5 RIC-3 knockdown reduces _α_7 and 5-HT_3A_ levels in SH-SY5Y plasma membrane fractions

GAPDH knockdown (GAPDH KD) in SH-SY5Y cells was achieved using 5 nM Silencer GAPDH siRNA at 50% confluence resulting in ∼95% reduction after 72 hours (**Fig. 3A**). We next optimized RIC-3 knockdown for use in a pull-down assay (PDA) with L1-MX and compared siRNA-transfected cells to non-transfected SH-SY5Y controls. In Fig. 3B, the left blot shows a clear RIC-3 band (∼50 kDa band, lane 9) in non-transfected ER fractions and only a faint band in RIC-3 siRNA samples (RIC-3 KD, lane 10). The membrane was then reprobed for calnexin (right blot), which remained unchanged after normalization, confirming comparable ER input across samples. Together, these data show that RIC-3 levels were reduced by >80% in transfected cells (**Fig. 3B**).

To assess whether RIC-3 knockdown impacted the surface expression of nAChα7 and 5-HT_3A_ receptors (both ∼50 kDa), we analyzed PM fractions from untransfected (RIC-3) and RIC-3 KD SH-SY5Y cells for α7 and 5-HT_3A_ expression (**Fig. 3C-D**). Blots were reprobed and normalized to Na^+^ /K^+^-ATPase. RIC-3 knockdown markedly reduced both receptors in the PM fraction, with quantification showing ∼98% reduction in 5-HT_3A_ and α7 compared to controls.

### 3.4 RIC-3 interacts with the 5-HT_3A_ L1-MX domain

To investigate RIC-3 interaction with 5-HT_3A_ ICD, we performed a PDA using the synthetic L1-MX peptide (25) (7) (34) (26). Consistent with earlier findings, RIC-3 from *X. laevis* oocytes interacted with the 5-HT_3A_ ICD (**Fig. 4B**) (25). The optimized assay was then applied to mouse brain and SH-SY5Y cell fractions, with bound proteins eluted, separated by SDS-PAGE, and probed by Western blot.

In *X. laevis* oocytes injected with RIC-3 cRNA, solubilized membrane preparations showed robust interaction between RIC-3 and the L1-MX–modified resin, with a clear 50□kDa band corresponding to monomeric RIC-3 (**Fig. 4B**, lane 4). The band intensity was approximately twice as strong compared to control resin capped with cysteine (Cys-resin). A similar enrichment pattern was observed in mouse brain ER fractions, where L1-MX resin captured more than twofold higher levels of RIC-3 than the capped control, with a prominent ∼50□kDa band and a faint ∼25□kDa band, potentially representing a processed or fragmented form (**Fig. 4C**, lane 7).

In contrast, SH-SY5Y ER fractions exhibited a different pattern: L1-MX resin predominantly pulled down RIC-3 at ∼100□kDa, with only faint or undetectable ∼50□kDa signal, suggesting the presence of higher-order oligomers (**Fig. 4D**, lanes 5-6). In these samples, enrichment over control was even greater, reaching approximately threefold. To determine whether the RIC-3 observed in the cytosolic fraction could also participate in this interaction, we subjected SH-SY5Y cytosolic lysates to the same pull-down conditions. Although Western blotting had previously revealed RIC-3 signals in the cytosol (**Fig. 2D**, lane 5), no RIC-3 band was recovered from the cytosolic eluate, indicating that this pool does not stably interact with the synthetic 5-HT_3A_ peptide under PDA conditions (**Fig. 4D**, lanes 7-8).

The key difference among oocytes, mouse ER, and SH-SY5Y samples was the relative abundance of extractable RIC-3, with SH-SY5Y cells containing the lowest levels and requiring more sensitive detection. The ∼100 kDa band in these cells was particularly sensitive to β 2-mercaptoethanol concentration and heating, necessitating gentler elution conditions for efficient recovery. RIC-3 at ∼50 kDa corresponds to the expected monomeric form, whereas the ∼100 kDa band suggests higher-order oligomers. Importantly, no interaction was detected in the RIC-3 knockdown sample (5 nM siRNA), where only nonspecific binding bands were observed (**Fig. 4E**).

Lanes with Cys-capped resin lacking L1-MX produced only faint nonspecific signals, confirming interaction specificity. Band intensities were quantified using Bio-Rad ImageLab software and normalized to the L1-MX–RIC-3 signal (100%), with Cys-capped controls (Cys-resin) showing >50% reduced binding across all conditions. Statistical analysis using GraphPad Prism (unpaired t-test), confirmed significant differences (p□<□0.05, **Fig. 4B-E**).

### 3.5 RIC-3 deglycosylation and migration in ER and PM fractions

To assess RIC-3 glycosylation, ER fractions were treated with PNGase F or Endo H under non-denaturing conditions. As shown in Fig. 5, the ER fraction exhibited a characteristic double band of RIC-3. Treatment with PNGase F removed the upper RIC-3 band, confirming N-linked glycan cleavage, while Endo H had no effect, indicating complex, Endo H-resistant glycosylation of RIC-3 (**Fig. 5A**). Fig. 5B further illustrates the differential migration of RIC-3 in ER and PM fractions when resolved on 4–20% gradient and 10% polyacrylamide gels. In the ER, a distinct double band of RIC-3 was observed in both gel types. However, the PM fraction showed only a prominent upper band on the gradient gel and up to six faint bands on the 10% gel, indicating a more heterogeneous and likely mature glycosylation profile in the PM.

**Fig. 5.**
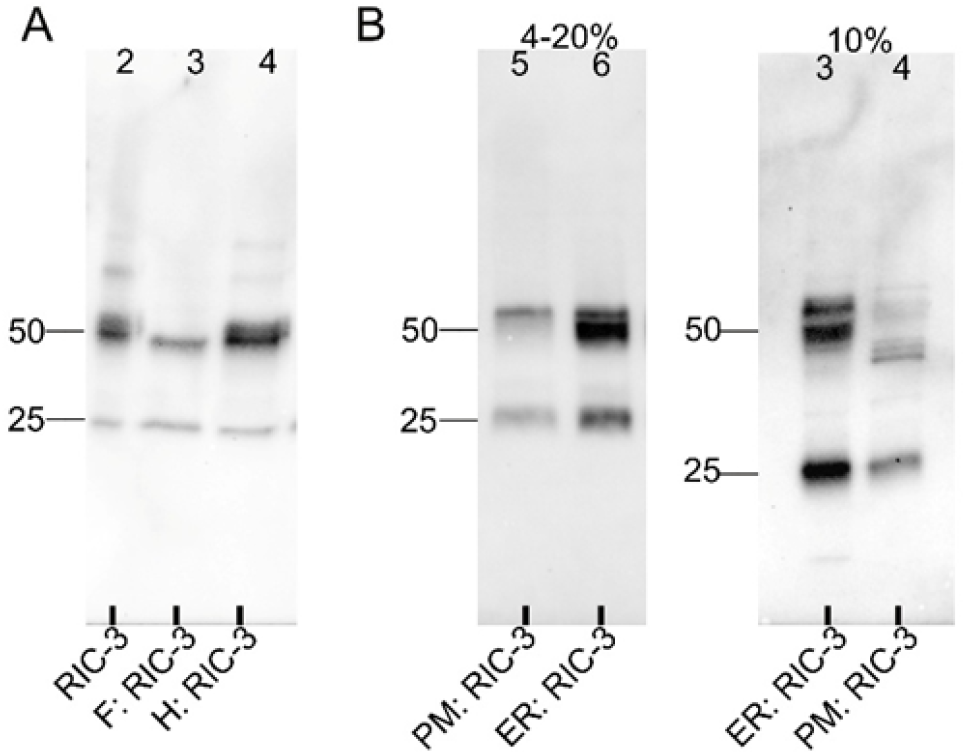
Glycosylation of RIC-3. **A** A double band of RIC-3 (50 kDa; lane 2) is observed in the mouse ER fraction. This fraction was subjected to deglycosylation using PNGase F (F: RIC-3) and Endo H (H: RIC-3) under non-denaturing conditions. PNGase F treatment resulted in the disappearance of the upper RIC-3 band (lane 3), indicating the removal of N-linked glycans. Endo H had no effect on RIC-3 (lane 4), suggesting resistance to Endo H-sensitive glycan cleavage. **B** Comparison of mouse RIC-3 electrophoretic profiles in ER and PM fractions. In both the gradient 4-20% (blot on the right, lane 6) and 10% gels (blot on the left, lane 3), ER-localized RIC-3 appears as characteristic upper and lower bands (∼50 kDa). In contrast, the PM fraction shows a prominent upper band with little to no lower band on the gradient gel (∼50 kDa, lane 5). When resolved on a 10% gel, the PM fraction displays multiple faint bands (lane 4). The ∼25 kDa bands present in both fractions likely represent a processed or fragmented form of RIC-3. Blots in Fig. 5 were cropped for clarity and presentation purposes. Molecular-weight marker positions (kDa) are indicated on the sides of each blot.

## 4. Discussion

RIC-3 is known to regulate the maturation and trafficking of cation-selective Cys-loop receptors (e.g., nAch, 5HT3) (20) (21) yet the structural basis of these interactions is still only partially defined. In our earlier work, we identified a duplicated RIC-3-5-HT_3A_ binding motif, but its endogenous nature and precise subcellular localization remained unresolved. Clarifying these features is essential for understanding how RIC-3 engages its receptor partners during biogenesis.

Our results here support the longstanding hypothesis that RIC-3 interacts with the ICD of the 5-HT_3A_ receptor, as demonstrated across *Xenopus laevis* oocytes, mouse brain tissue, and neuroblastoma cell lines (**Fig. 4B-E**). We confirmed the cellular distribution of RIC-3 and identified the subcellular potential sites where these interactions are likely to occur (**Fig.s 2C-D, 4B-E, 5B**). RIC-3 is primarily recognized as an ER-resident chaperone involved in the maturation and trafficking of nAChRs, particularly α7 subunits (35). Consistent with this, mouse brain fractionation revealed that RIC-3 was predominantly localized to the ER, with only a faint band in the crude PM fraction and two faint ∼50 kDa bands in the cytosol (**Fig. 2C**). While RIC-3 is not typically abundant at the plasma membrane, the faint band detected in the PM fraction suggests that a small pool may localize to this region, potentially reflecting either a transient trafficking state or a functional subpopulation (**Fig. 5B**) (25). This band could alternatively arise from membrane-associated ER compartments (36) (29). Similarly, the weak RIC-3 signal observed in mouse cytosol may reflect minor cross-contamination during fractionation (**Fig. 2C**) (29). In contrast, in SH-SY5Y cells, fractionation consistently revealed a weaker RIC-3 signal in the cytosolic fraction relative to the ER (**Fig. 2D**, lane 5-6). However, pull-down assays from cytosolic lysates did not recover RIC-3 (**Fig. 4D**, lane 7-8). This discrepancy indicates that the SH-SY5Y cytosolic signal likely represents low-abundance or unstable RIC-3 species, either fragments targeted for degradation or minor ER carryover, rather than a functional pool capable of stable interactions. Thus, the ER fraction appears to be the primary compartment from which RIC-3 can be effectively enriched in PDA. As expected, no RIC-3 signal was detected in mitochondrial fractions (**Fig. 2 C-D**).

The RIC-3 protein is known to exist in multiple isoforms, reflecting its diversity through alternative mRNA splicing patterns (37). Although alternative splicing is believed to enhance protein diversity from a fixed genome, there is ongoing debate about whether many of these splice variants are truly functional or simply the result of splicing errors (38). In the mouse brain ER fraction, RIC-3 appears as two closely spaced bands around 50 kDa, and this doublet pattern was consistently observed in both gradient and 10% SDS-PAGE gels (**Fig. 2C**, lane 6, **5B**). Treatment with PNGase F removed the upper band, indicating that it was caused by N-linked glycosylation (**Fig. 5A**, lane 3). A similar result was previously observed using Endo H, which also removes certain types of N-linked sugars, although it targets a narrower range than PNGase F (33). This indicates that the core RIC-3 protein has a MW of ∼44 kDa, while a subset undergoes extensive glycosylation, increasing its apparent MW to ∼50 kDa. Our data show that only the upper glycosylated form of RIC-3 was consistently detected in the PM fraction (**Fig. 5B**, lane 5). When resolved on 10% gels (lane 4), the PM sample displayed multiple faint bands, indicating increased heterogeneity and further supporting the hypothesis that RIC-3 undergoes additional processing during ER-to-PM transit. Although canonically described as an ER-resident chaperone for 5-HT_3A_ receptor assembly, our findings suggest that a subset of RIC-3 may escape ER retention and accompany its client receptor through the secretory pathway, potentially undergoing a core-to-complex glycosylation transition and reaching the cell surface via the Golgi apparatus. We propose that the presence of glycosylated RIC-3 at the plasma membrane reflects a regulated, physiologically relevant event, either representing a transitional phase during receptor complex export or a sustained chaperone function beyond the ER. While one possibility is that RIC-3 reaches the surface due to incomplete dissociation from its client receptor, the consistent detection of a specific glycoform at the PM and its distinct localization pattern argue instead for a deliberate, functional extension of its role. Surface-localized RIC-3 may contribute to late-stage receptor maturation, membrane stabilization, or glycan-mediated extracellular interactions.

RIC-3 plays a key role in helping nAChRs and 5-HT_3A_ receptors function properly by supporting the assembly of their subunits into active receptors on cell membranes. A recent study investigated how RIC-3 impacted nAChR expression in the ER (39). These findings suggest that RIC-3 could be a promising target for treating neurodegenerative diseases and other conditions involving disrupted cholinergic signaling. The study also suggests the potential of RIC-3-dependent therapies for tau-related disorders, highlighting RIC-3’s importance in brain function and future treatment approaches.

In our previous study, we established an interaction between the MBP-5-HT_3A_-ICD construct and RIC-3 (26) (27) (25). This aligns with the results of the current study, which found that RIC-3 from the PM/ER fraction of three sources (frog oocytes/mouse brain/cell lines) displayed interactions with the synthetic 5-HT_3A_ peptide. We observed a ∼50 kDa band corresponding to monomeric RIC-3, whereas the additional ∼100 kDa species suggests oligomerization and possible complex assembly. Elaborating on these findings, interactions between serotonin receptors and RIC-3 suggests a potential regulatory role for RIC-3 in influencing receptor function and expression.

In this study, we achieved more than 80% knockdown of RIC-3 in SH-SY5Y cells after 24-hour incubation with siRNA (**Fig. 3B**). This reduction lowered RIC-3’s interaction with the synthetic L1-MX peptide (**Fig. 4E**, lane 8) and sharply reduced surface expression of α7 nAChRs and 5-HT_3A_ receptors (**Fig. 3C-D**), supporting our hypothesis that RIC-3 is critical for receptor interactions and trafficking. Reduced receptor availability at the plasma membrane is likely to impair synaptic transmission and contribute to excitatory/inhibitory imbalances relevant to psychiatric disorders (4). Altered α7 nAChR expression is well documented in schizophrenia, where it contributes to sensory gating and cognitive deficits; clinical trials showed that α7 agonists can improve cognitive performance and reduce negative symptoms (40). Although direct evidence for reduced 5-HT_3A_ expression in patients is limited, its dysregulation has been linked to anxiety and depression (2), autism spectrum traits (41), and explored as a therapeutic target in schizophrenia (42). Our findings reflect a transient 24-hour knockdown of RIC-3; however, with prolonged or chronic deficiency, persistent ER stress would be expected to cause protein buildup, neuronal dysfunction, or cell death. Although cells may initially compensate by upregulating ER chaperones such as calnexin, calreticulin, and BiP, this response may not fully restore homeostasis. Consistent with this, transient knockdown did not change calnexin levels (**Fig. 3B**). With prolonged loss or complete knockout, compensatory mechanisms would likely fail, leading to protein aggregation, chaperone imbalance, and widespread dysfunction.

Beyond psychiatry, RIC-3 dysfunction has been linked to Parkinson’s disease (43), multiple sclerosis (44), and cholinergic dysregulation implicated in Alzheimer’s disease. Together, these findings position RIC-3 as a critical regulator of receptor maturation, integrating cholinergic, serotonergic, dopaminergic, and immune signaling, with therapeutic potential for neuropsychiatric, neurodegenerative, and neuroinflammatory diseases (45).

## Contributions

MJ and NJ designed research. MJ and NJ performed research. Petar N. Grozdanov provided mouse brain tissue and assisted with three different techniques. Clinton C. MacDonald supplied mouse brain tissue. Rhea Ramani, Cade Perkins, Hoa Q Do, and Joshua Theriot assisted with experimental preparations. NJ wrote the paper and all authors read and approved the final manuscript.

**Note:** Nermina Jahovic and Nermina Sarayli Belirgen are one and the same individual. The name, Nermina Jahovic, represents the scientist’s maiden name, and this statement has been written at the scientist’s request.

## Acknowledgement

We thank the TTUHSC Core Facilities; some of the images and/or data were generated in the Image Analysis Core Facility and Molecular Biology Core Facility supported by TTUHSC. Research reported in this publication was supported by the National Institute of Neurological Disorders and Stroke (NINDS) of the National Institutes of Health (NIH) under award number NS077114 (to M.J.).

